# Fast Bayesian fine-mapping of 35 production, reproduction and body conformation traits in dairy cattle

**DOI:** 10.1101/428227

**Authors:** Jicai Jiang, John B. Cole, Yang Da, Paul M. VanRaden, Li Ma

## Abstract

Imputation has been routinely used to infer sequence variants in large genotyped populations based on reference populations of sequenced individuals. With increasing numbers of animals sequenced and the implementation of the 1000 Bull Genomes Project, fine-mapping of causal variants for complex traits is becoming possible in cattle. Using 404 ancestor bull sequences as reference, we imputed over 3 million selected sequence variants to 27,214 Holstein bulls with highly reliable phenotypes (breeding values) for 35 production, reproduction, and body conformation traits. We first performed whole-genome single-marker scans for each of the 35 traits using a mixed-model association test. The single-trait association statistics were then merged into multi-trait tests of 3 trait groups: production, reproduction, and body conformation. Both single- and multi-trait GWAS results were used to identify 282 candidate QTL regions for fine-mapping in the cattle genome. To facilitate fast and powerful fine-mapping analyses, we developed a Bayesian Fine-MAPping approach (BFMAP) to integrate fine-mapping with functional enrichment analysis. Our fine-mapping results identified 69 promising candidate genes for dairy traits, including *ABCC9, VPS13B, MGST1, SCD, MKL1*, and *CSN1S1* for production traits; *CHEK2, GC*, and *KALRN* for reproduction traits; and *TMTC2, ARRDC3, ZNF613, CCND2*, and *FGF6* for body conformation traits. Based on existing functional annotation data for cattle, we revealed biologically meaningful enrichment in our fine-mapped variants that can be readily tested in functional validation studies. In summary, these results demonstrated the utility of a fast Bayesian approach for fine-mapping and functional enrichment analysis, identified candidate causative genes and variants, and enhanced our understanding of the genetic basis of complex traits in dairy cattle.

## Introduction

Phenotypic records have been routinely collected in dairy cattle to facilitate selective breeding for more than one hundred years. The phenotype of a bull can be highly accurately calculated from thousands of phenotypic records of his daughters and other relatives^1^. A comprehensive spectrum of phenotypes has been recorded in dairy cattle, including production, reproduction, health, and body type traits ^2^. GWAS on these traits simultaneously in the same population can provide a better understanding of the effects of underlying QTLs. Because of the intensive use of artificial insemination and strong selection in dairy bulls, there are a much smaller number of males than females in the cattle population ^3^, and chromosome segments can be quickly traced back to an ancestral bull. The high relatedness in the cattle population can facilitate accurate imputation ^4^, especially with many important ancestor bulls sequenced by the 1000 Bull Genomes project ^5^. These unique features of the cattle population make a large-scale GWAS with imputed sequence variants possible and valuable.

Fine-mapping of complex traits to single-variant resolution has been possible in human studies, e.g., ^6,7^. Because of the high levels of linkage disequilibrium (LD) in the cattle population ^8^, fine-mapping of GWAS signals has been difficult. In addition, existing fine-mapping methods are not readily applicable to large-scale cattle GWAS and fine-mapping studies. These methods, e.g., CAVIARBF ^9^ and PAINTOR ^10^, generally use a logistic model with a binary response and categorical functional annotations as covariates. Such a logistic model is then incorporated into a model search scheme that often limits the maximum number of causal variants (e.g., 3) and is computationally impractical for a locus containing thousands of sequence variants. When multiple functional data sets are to be tested, model-searching needs to be conducted separately for each set of functional annotation data, further increasing the computational burden. To address these problems, we propose a fast Bayesian Fine-MAPping method (BFMAP) that can efficiently integrate functional annotations with fine-mapping. Specifically, BFMAP can re-use initial model search results for various functional annotations and can be employed for both fine-mapping and functional enrichment analyses. More importantly, the functional enrichment estimated from BFMAP is, by definition, the *enrichment of causal effects*, in contrast to the *enrichment of heritability* by the well-known stratified LD score regression ^11^.

In our study, the large number of bulls with highly reliable phenotype and imputed sequence variants can facilitate powerful GWAS and fine-mapping of major GWAS signals. Although the high LD in the cattle genome makes fine-mapping and functional enrichment studies difficult, the large sample size and improved methods can help identify candidate genes of complex traits as well as biologically informative enrichment of candidate variants in functional annotation data. Specifically, we seek to use BFMAP to identify and incorporate functional annotation into the fine-mapping of 35 production, reproduction, and conformation traits in dairy cattle. The fine-mapped genes and variants can provide candidates readily testable in functional studies. The functional data enriched with variants associated with complex dairy traits will be useful for future cattle GWAS and genomic prediction studies. Additionally, the initial model search results can be reused for estimating enrichment of causal effects of dairy traits for additional functional annotations that are being generated by the FAANG and related projects in cattle ^12^.

## Results

We imputed over 3 million selected sequence variants to 27,214 Holstein bulls after quality control edits, using the 1000 Bull Genomes data as reference. These bulls were selected to have highly reliable breeding values (PTA) for 35 production, reproduction, and body conformation traits, with an average reliability of 0.71 across traits (Table 1). The numbers of bulls available for individual traits ranged from 11,713 to 27,161, with >20,000 animals having data for 32 traits (Table 1). This large, high-quality bull data set enables our following GWAS and fine-mapping studies with great power and precision.

**Table 1.** Number of Holstein bulls, and mean and standard deviation (SD) of PTAs and reliabilities for 35 dairy traits.

| Trait Name | Abbreviation | N of Bulls | Deregressed PTA |  | Reliability |  |
| --- | --- | --- | --- | --- | --- | --- |
|  |  |  | Mean | SD | Mean | SD |
| Milk yield | Milk | 27156 | -245.86 | 850.58 | 0.860 | 0.082 |
| Fat yield | Fat | 27156 | -5.92 | 30.52 | 0.860 | 0.082 |
| Protein yield | Protein | 27156 | -5.31 | 23.84 | 0.863 | 0.083 |
| Fat percentage | Fat_Percent | 27156 | 0.0136 | 0.107 | 0.860 | 0.082 |
| Protein percentage | Pro_Percent | 27156 | 0.0086 | 0.0464 | 0.863 | 0.083 |
| Net merit | Net_Merit | 27161 | -106.91 | 278.63 | 0.763 | 0.110 |
| Productive life | Prod_Life | 26727 | -1.367 | 3.461 | 0.682 | 0.145 |
| Somatic cell score | SCS | 27143 | 3.027 | 0.235 | 0.786 | 0.110 |
| Age at first calving | AFC | 16314 | -0.446 | 11.855 | 0.439 | 0.258 |
| Days to first breeding<sup>1</sup> | DFB | 11713 | 0.534 | 2.825 | NA | NA |
| Daughter pregnancy rate | Dtr_Preg_Rate | 25699 | -0.593 | 3.025 | 0.618 | 0.185 |
| Heifer conception rate | Heifer_Conc_Rate | 19334 | -0.660 | 9.610 | 0.377 | 0.210 |
| Cow conception rate | Cow_Conc_Rate | 20380 | -1.053 | 6.879 | 0.597 | 0.202 |
| Sire calving ease | Sire_Calv_Ease | 26345 | 7.959 | 2.461 | 0.671 | 0.224 |
| Daughter calving ease | Dtr_Calv_Ease | 23263 | 9.141 | 3.182 | 0.594 | 0.176 |
| Sire stillbirth | Sire_Still_Birth | 21543 | 8.190 | 1.831 | 0.495 | 0.249 |
| Daughter stillbirth | Dtr_Still_Birth | 20424 | 8.085 | 2.958 | 0.508 | 0.222 |
| Final score | Final_score | 25638 | -0.817 | 1.484 | 0.702 | 0.140 |
| Stature | Stature | 25641 | -0.482 | 1.532 | 0.844 | 0.079 |
| Strength | Strength | 25633 | -0.278 | 1.513 | 0.743 | 0.147 |
| Dairy form | Dairy_form | 25615 | -0.492 | 1.745 | 0.752 | 0.132 |
| Foot angle | Foot_angle | 25626 | -0.742 | 2.263 | 0.664 | 0.198 |
| Rear legs (side view) | Rear_legs(side) | 25641 | -0.009 | 1.734 | 0.754 | 0.137 |
| Body depth | Body_depth | 25636 | -0.413 | 1.622 | 0.720 | 0.180 |
| Rump angle | Rump_angle | 25641 | 0.038 | 1.482 | 0.828 | 0.089 |
| Rump width | Rump_width | 25641 | -0.504 | 1.543 | 0.766 | 0.114 |
| Fore udder attachment | Fore_udder_att | 25640 | -0.908 | 1.852 | 0.781 | 0.112 |
| Rear udder height | Rear_ud_height | 25640 | -0.885 | 2.095 | 0.737 | 0.136 |
| Udder depth | Udder_depth | 25631 | -0.653 | 1.665 | 0.836 | 0.082 |
| Udder cleft | Udder_cleft | 25641 | -0.720 | 1.980 | 0.718 | 0.156 |
| Front teat placement | Front_teat_pla | 25641 | -0.562 | 1.663 | 0.781 | 0.106 |
| Teat length | Teat_length | 25631 | 0.104 | 1.482 | 0.815 | 0.087 |
| Rear legs (rear view) | Rear_legs(rear) | 24763 | -0.759 | 2.709 | 0.605 | 0.178 |
| Feet and legs composite | Feet_and_legs | 25608 | -0.928 | 2.501 | 0.600 | 0.208 |
| Rear teat placement | Rear_teat_pla | 25492 | -0.436 | 1.900 | 0.762 | 0.103 |

<sup>1</sup>For DFB, we used PTA because reliability was unavailable.

### Single-trait GWAS

We used a mixed model approach implemented in the software MMAP^13^ that can incorporate reliability variation across individual bulls. The mixed model used in our GWAS was robust against population structure and familial relatedness. As shown in Table S1, 27 of the 35 traits had a genomic control factor between 0.95 and 1.05.

Using a genome-wide significance level of *P* < 5E-8, we found many clear association signals for the 35 dairy traits (Fig S1). There were in total 286 QTL regions associated with the 35 traits, and the number of associations for individual traits ranged from <3 for leg and foot traits to 23 for protein percentage (Tables S1 and S2). As compared to the Cattle QTLdb release 35^14^, we found that 123 associations (43%) had been previously reported while 163 associations (57%) were newly discovered in this study. We identified 15 new association signals (out of 68) even for the five production traits that had been extensively studied, and 92 new associations (out of 125) for type traits that drew less attention in previous studies (Fig 1 and Table S2). While a proportion of these newly discovered QTLs were identified to be associated with new traits, these results demonstrated the superior power of our GWAS in dairy cattle.

**Figure 1.**
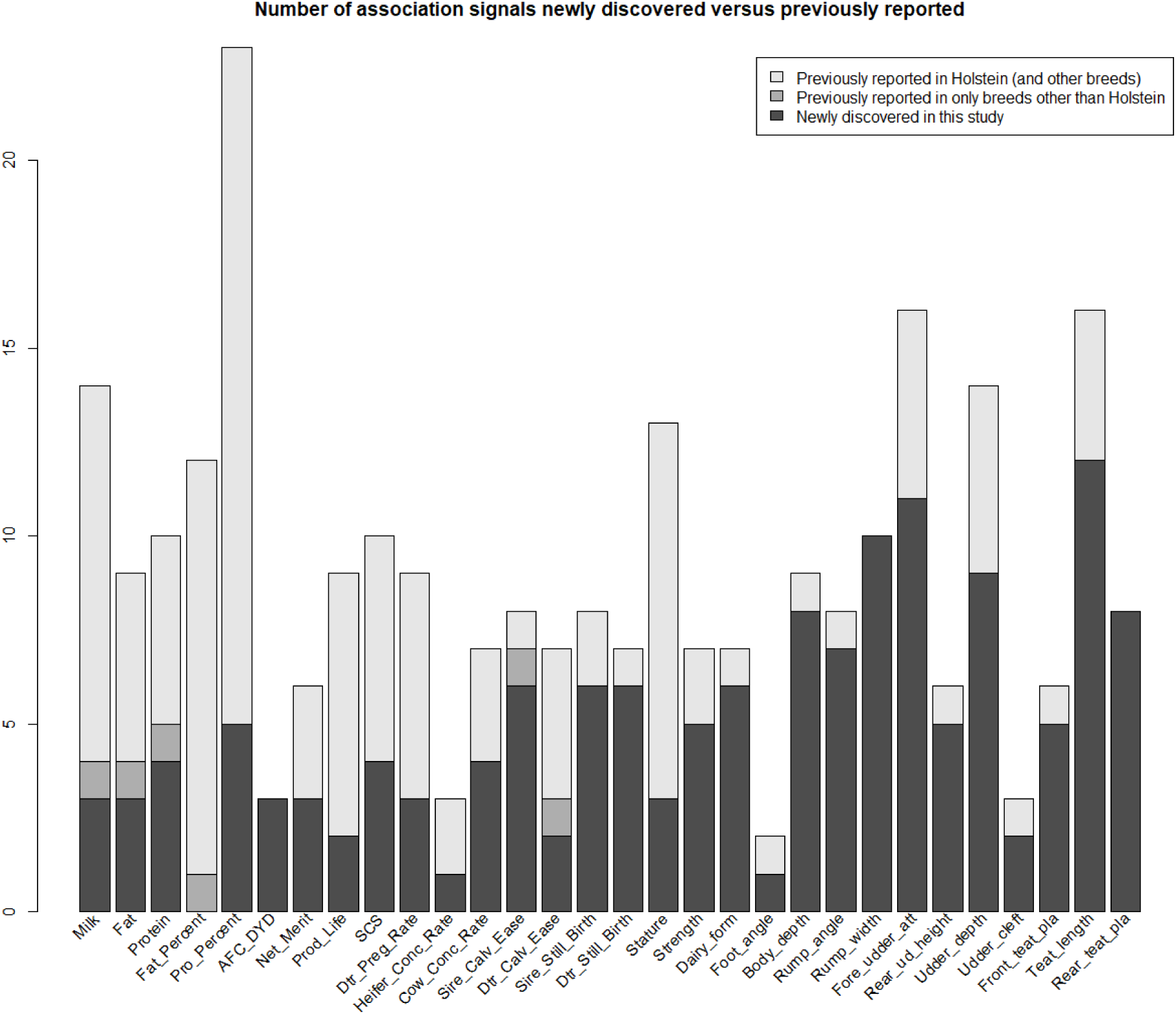
Number of association signals newly discovered in our single-trait GWAS versus previously reported. There are in total 30 traits listed. Three leg traits were excluded since we found no associations passing genome-wide significance. Days to first breeding (DFB) and final score were not listed because there were no matched traits in the Cattle QTLdb release 35.

### Multi-trait association analysis

Consistent with trait definition, hierarchical clustering of the 35 traits based on the absolute correlation coefficients identified three clusters: production, reproduction, and body type (Fig 2). Interestingly, rump angle and teat length were clustered into reproduction traits, although they are type traits by definition, indicating a close genetic correlation between these two traits and cattle reproduction.

**Figure 2.**
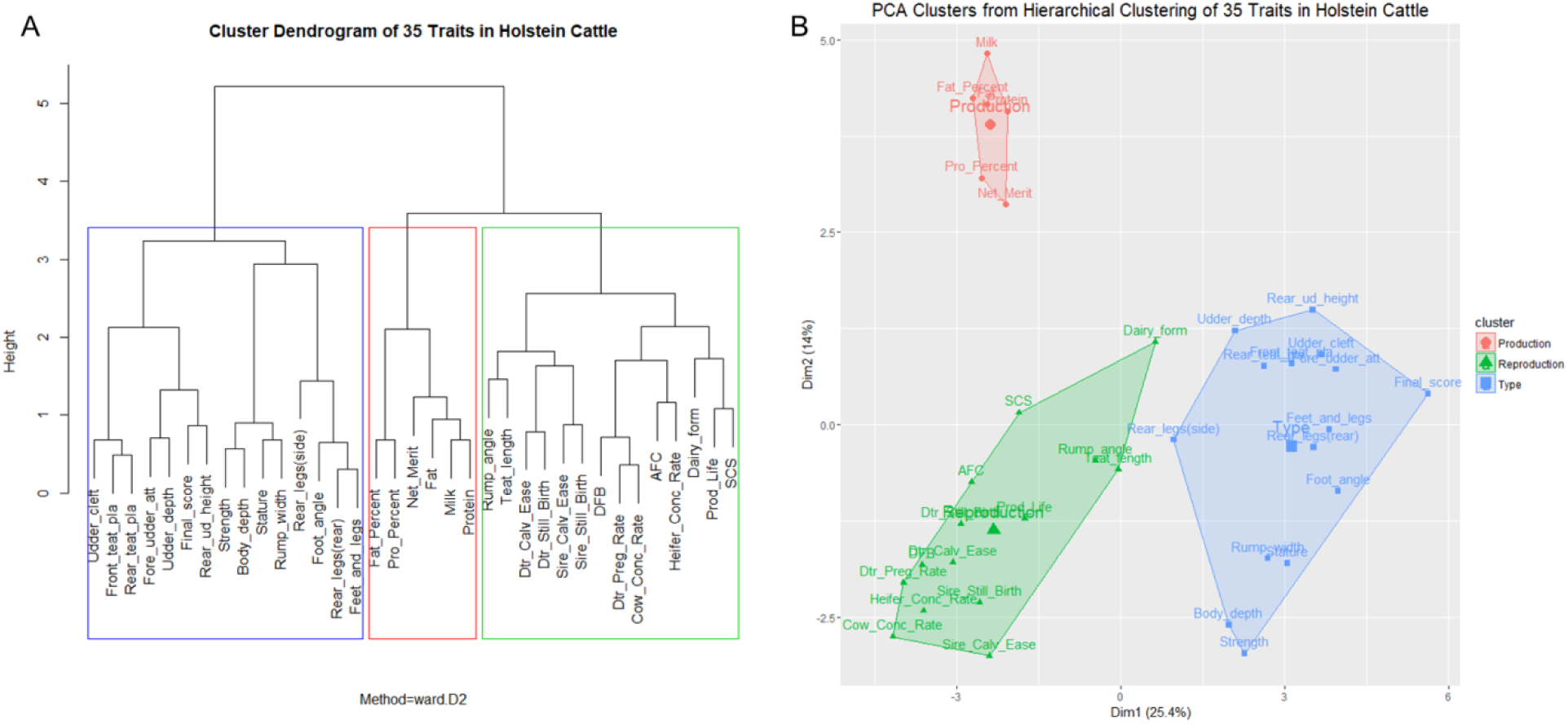
Hierarchical clustering of 35 traits in Holstein cattle. A. Cluster dendrogram. B. PCA clusters.

From multi-trait association analyses of the three trait clusters, we identified 33, 21, and 39 associations for production, reproduction, and type traits using *P* < 5E-8, respectively (Fig 3 and Table S3). Although the majority of the multi-trait associations were already identified from single-trait GWAS, we found ten associations that were missed by single-trait analyses (Table S4). Based on the multi-trait analysis results, we found two features of multi-trait association tests. First, multi-trait GWAS was more powerful than individual single-trait analyses for related traits. Second, the top variant in multi-trait analysis may be >1 Mb away from the top variants in single-trait GWAS (Fig S2).

**Figure 3.**
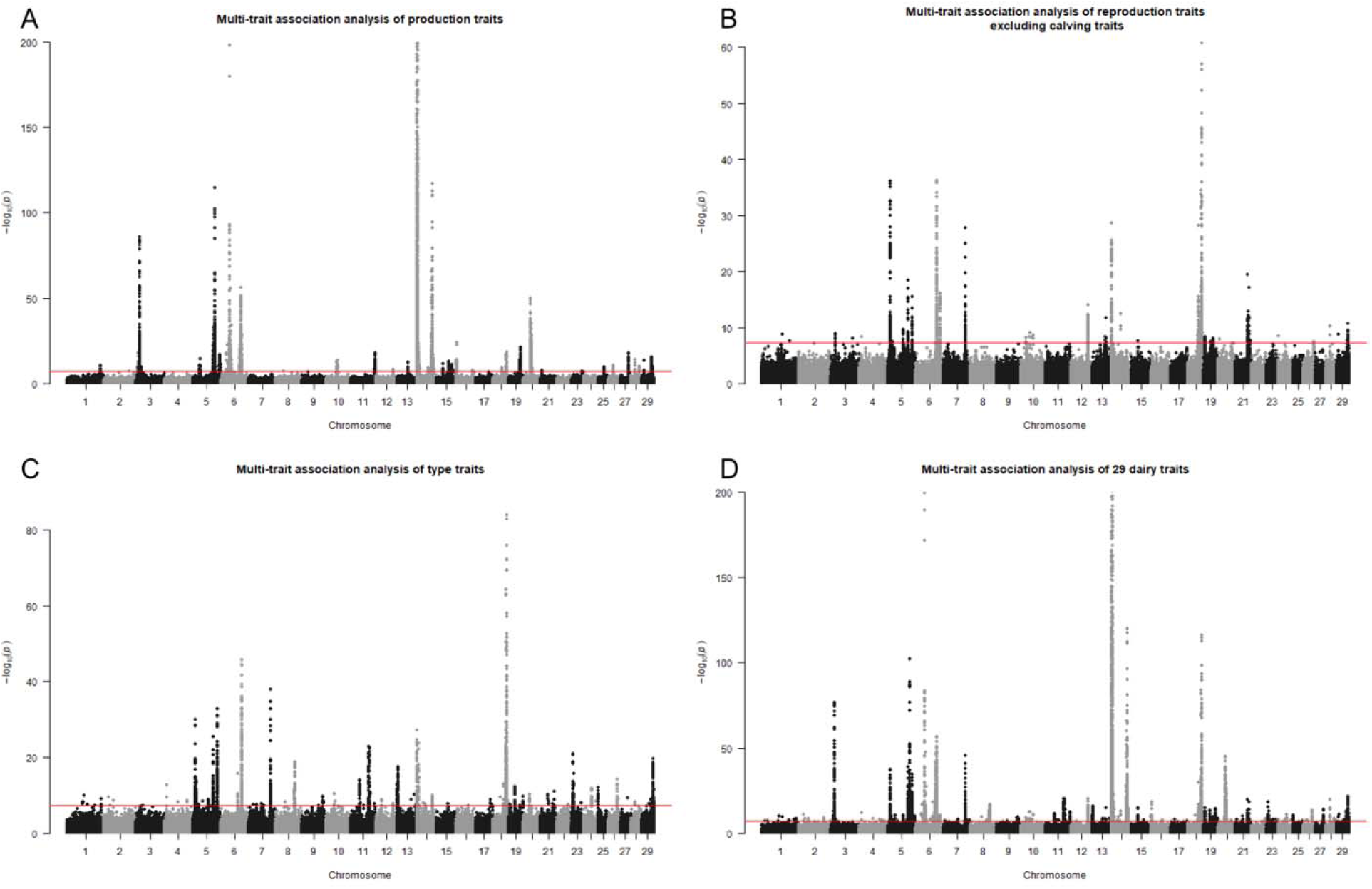
Manhattan plots for multi-trait association analyses. A. Production traits. B. Reproduction traits, excluding four calving traits (calving ease and stillbirth traits). C. Type traits. D. All 29 dairy traits, excluding days to first breeding (DFB), net merit, and four calving traits.

### Fine-mapping

To facilitate fast fine-mapping analyses, we developed a fast Bayesian Fine-MAPping method (BFMAP) that calculates a posterior probability of causality (PPC) for variants in candidate regions. We picked QTL regions for fine-mapping from both single- and multi-trait GWAS results. Initially, we fine-mapped 434 association signals for 282 QTLs using a significance threshold of 5E-7 (Table S5). The observed number of fine-mapped signals in a QTL is approximately exponentially distributed, consistent with our expectation of more causal mutations with a lower probability in a QTL region (Fig 4). After further quality control edits, we finally fine-mapped 308 association signals for 32 traits (Table S6). Specifically, there were more than 20 independent association signals identified on chromosomes 5, 6, 14, 18, and 29, while very few were identified on chromosomes 12, 22, and 27.

**Figure 4.**
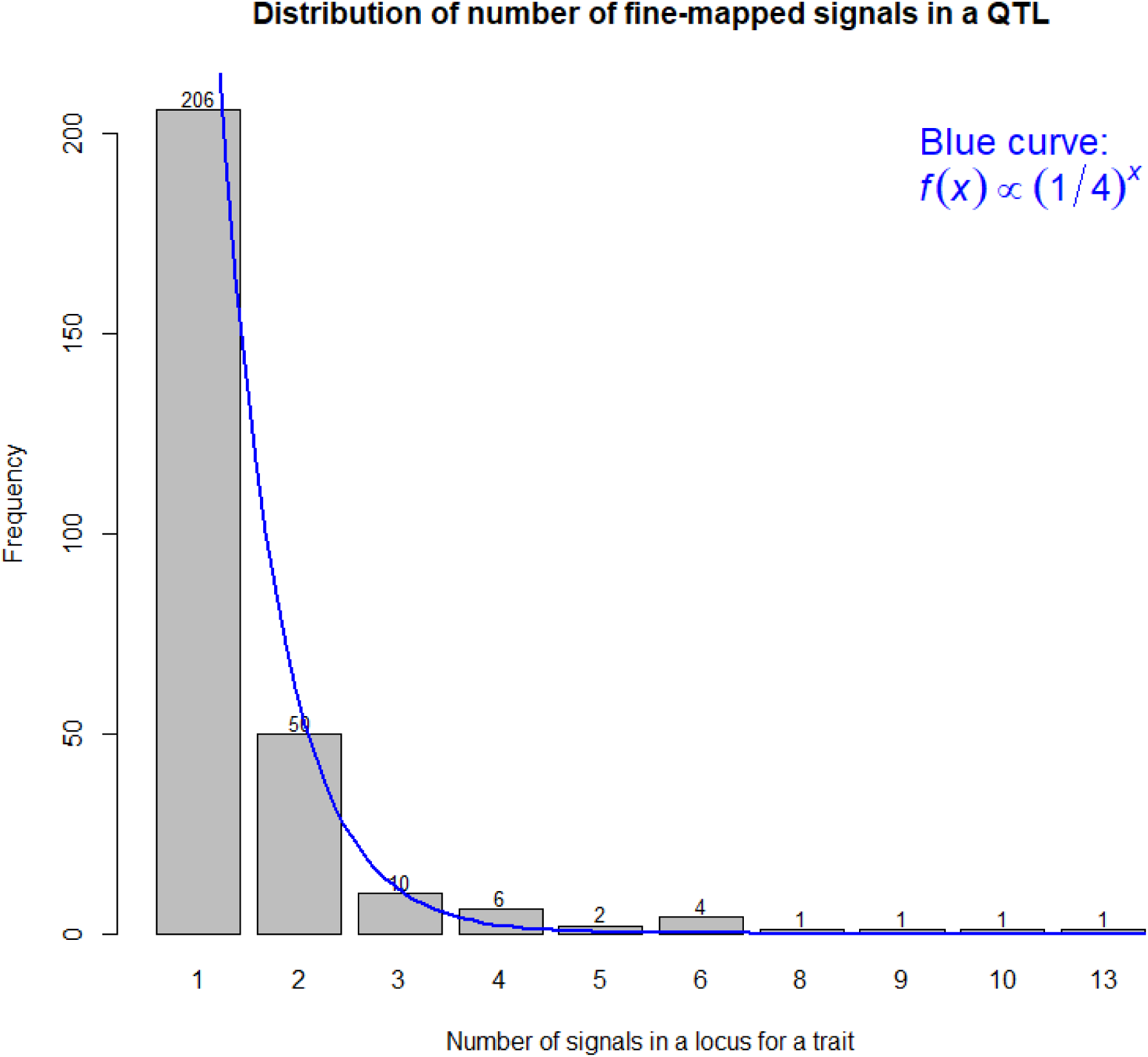
Distribution of number of fine-mapped signals in a candidate locus for a trait. Signals were filtered by a significance threshold of 5*E*-7.

We investigated the impacts of incorporation of SnpEff-inferred effect impact (commonly used functional annotation) on fine-mapping performance. First, incorporating variant impacts resulted in a substantial change of PPC for variants in the 308 fine-mapped association signals. Variants with moderate impact had a considerable increase in PPC when functional information was included in the calculation, while modifier variants generally had a decreased PPC (Fig 5A). Second, fine-mapping by incorporating variant impacts generated significantly smaller 95% credible variant sets than that using an equal prior for all variants (*P*<0.01, Wilcoxon signed-rank test; Fig 5B). These two features make the incorporation of functional annotation favored in our fine-mapping analyses.

**Figure 5.**
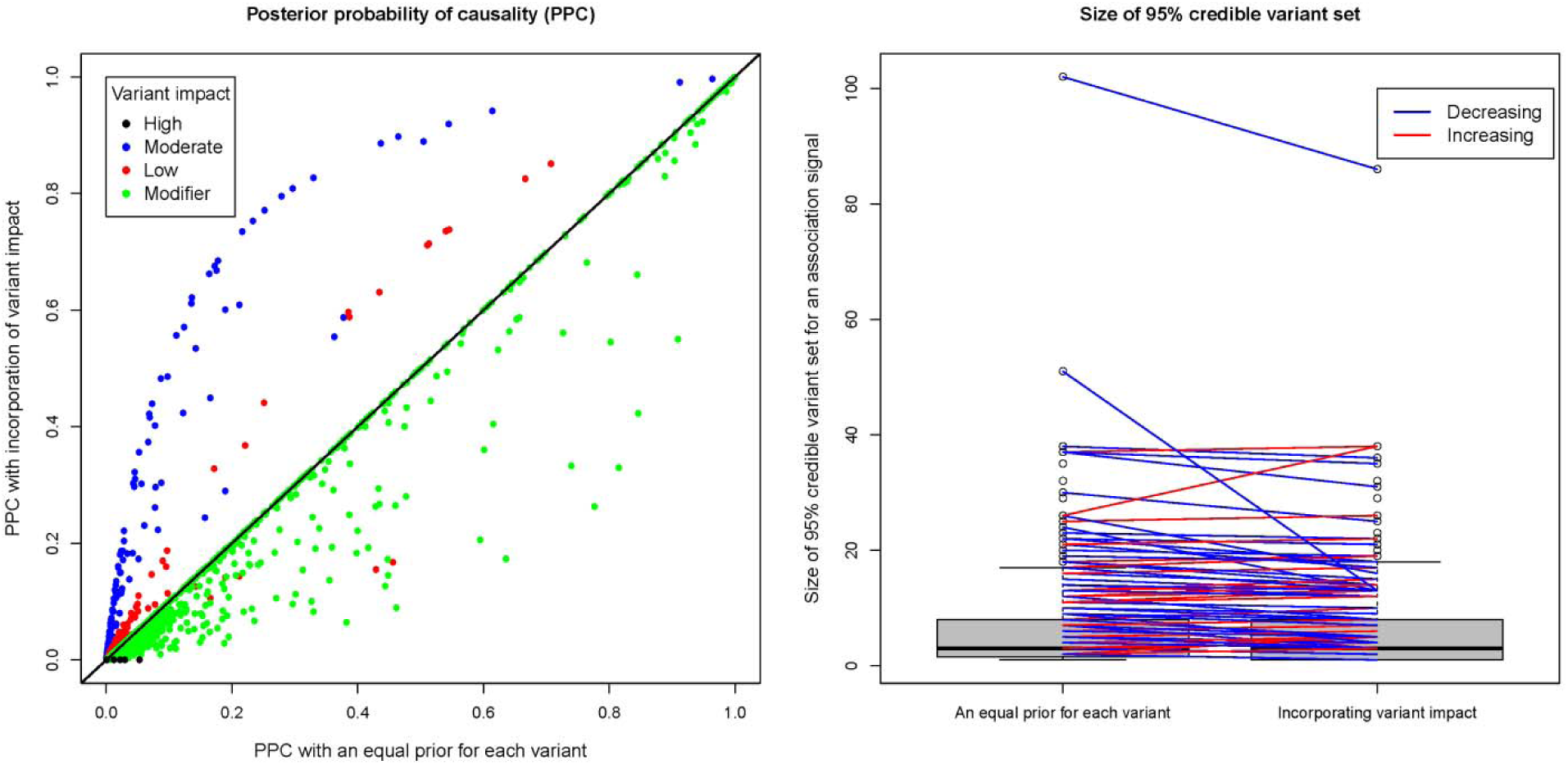
Effect of incorporation of SnpEff-inferred impact on fine-mapping performance. A. Posterior probability of causality (PPC) with incorporation of SnpEff impacts versus PPC with an equal prior for each variant. B. Size of 95% credible variant set generally decreased after incorporation of SnpEff-inferred impact.

### Enrichment analysis

To verify the quality of our fine-mapped variants and characterize their distribution on the cattle genome, we investigated the enrichment of fine-mapped variants with different functional annotation data available to cattle, including location in protein-coding gene, effect predicted by SnpEff ^15^, and evolutionary constrain predicted by GERP ^16^. Our enrichment analysis estimated the probability of a causal variant being in a functional category and the probability of a non-causal variant being in the category. The ratio of the two probabilities was used to measure the enrichment of causal variants for this functional category ^17^, with a value larger than one indicating higher enrichment than the genome background. This enrichment analysis has also been implemented in BFMAP.

We first categorized variants into five groups based on their locations regarding protein-coding genes, i.e., CDS, 5’ UTR + 2 kb upstream, intron, 3’ UTR + 2 kb downstream, and other (intergenic or non-protein-coding genic regions). Despite the strong LD levels in the cattle genome ^18^, we observed distinctive enrichment patterns across these five categories (Fig 6A). Using bootstrapping, we calculated 95% confidence intervals for the enrichment levels, showing significant enrichment of fine-mapped variants in CDS (4.52x) and 5’ UTR (2.39x), but not in intron (0.93x) or 3’ UTR (0.77x). We also analyzed a group of non-protein-coding genes but found significant depletion with 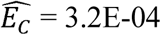 (Table S7), suggesting a lacking of functional impacts in these genes on dairy cattle traits.

**Figure 6.**
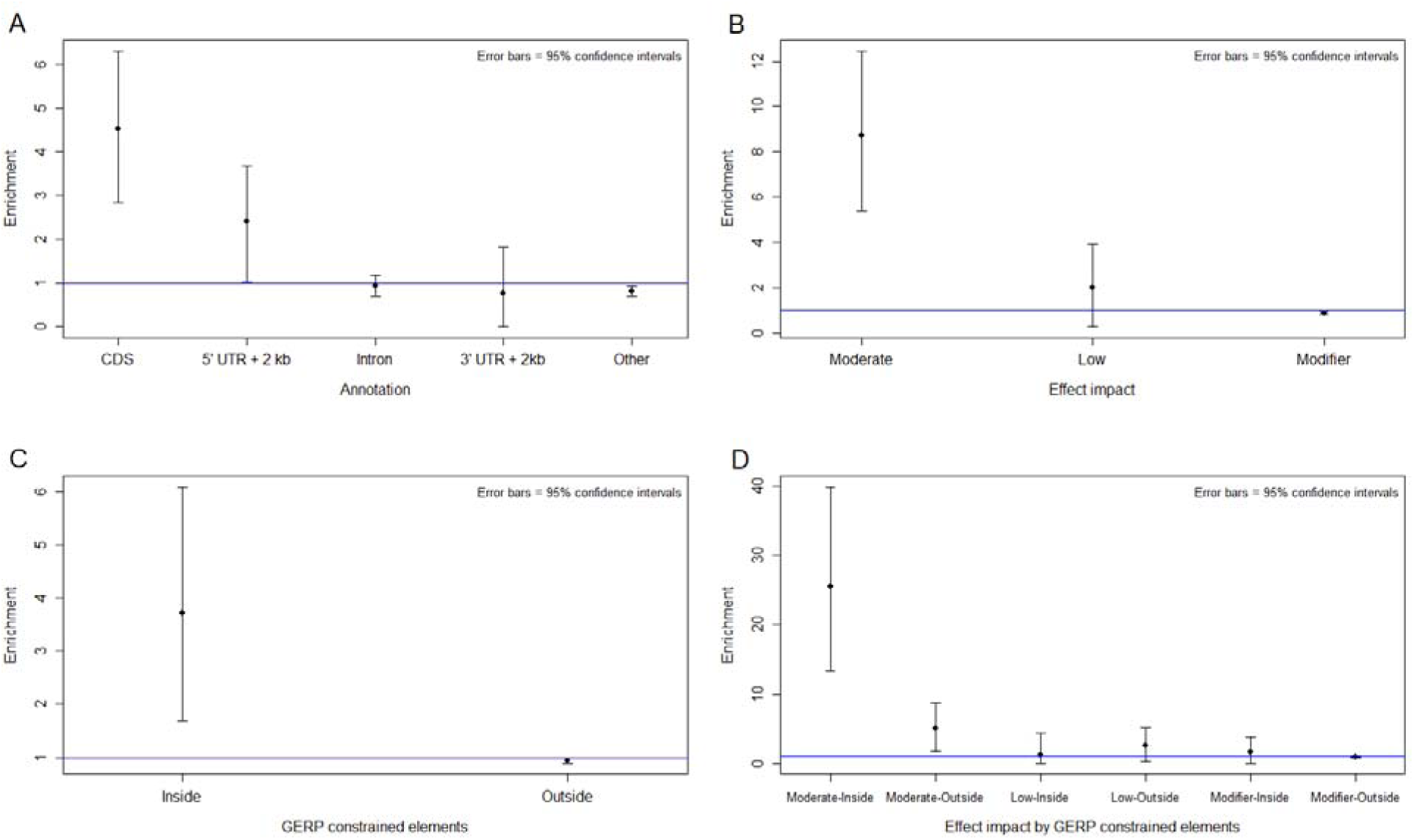
Enrichment of fine-mapped variants across various functional annotations. A. Locations of variants regarding protein-coding genes. B. SnpEff predicted impact. C. GERP-constrained elements. D. SnpEff impact and GERP-constrained elements.

We further investigated the enrichment of fine-mapped variants regarding their genomic locations and protein coding effects (High, Moderate, Low or Modifier) predicted by SnpEff ^15^. When modeling these four categories, we found severe depletion of variants with high impact (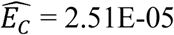; Table S8). This is strikingly different from a previous study on human complex traits and diseases that reported an enrichment of >100 for this category ^17^. As shown in Fig 5B, we observed a significant enrichment in moderate-impact variants (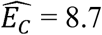; *P* < 0.05). Low-impact variants also showed an enrichment (2.0x), though it was not statistically significant (Fig 6B). As expected, a minor depletion was seen in modifier variants (0.87x).

We also used constrained elements on the cattle genome to categorize variants into two groups (inside of or outside of constrained elements), as highly conserved DNA sequences may imply functional importance. As shown in Fig 6C and Table S9, fine-mapped variants were significantly enriched in constrained elements (3.72x; *P* < 0.05). When further categorizing variants into six groups based on both constrained elements and variant impacts (Moderate, Low or Modifier), we found the highest enrichment in moderate-impact variants inside constrained elements (25.56x; *P* < 0.05). For the other categories, we observed no significant enrichment of fine-mapped variants (Fig 6D and Table S20).

When comparing different trait groups, we observed little difference in the pattern of enrichment regarding SnpEff-inferred effect impact (Fig 7 and Table S21). Moderate-impact variants had a clearly higher enrichment of being causal for production traits than for reproduction and type traits. We further used permutation to generate the null distribution of *E_C_*(Production)/*E_C_*(Reproduction+Type) and showed that the difference was statistically significant (*P* < 0.05; Fig S3A). However, the enrichment for low-impact variants was similar between the three trait groups (Fig S3B).

**Figure 7.**
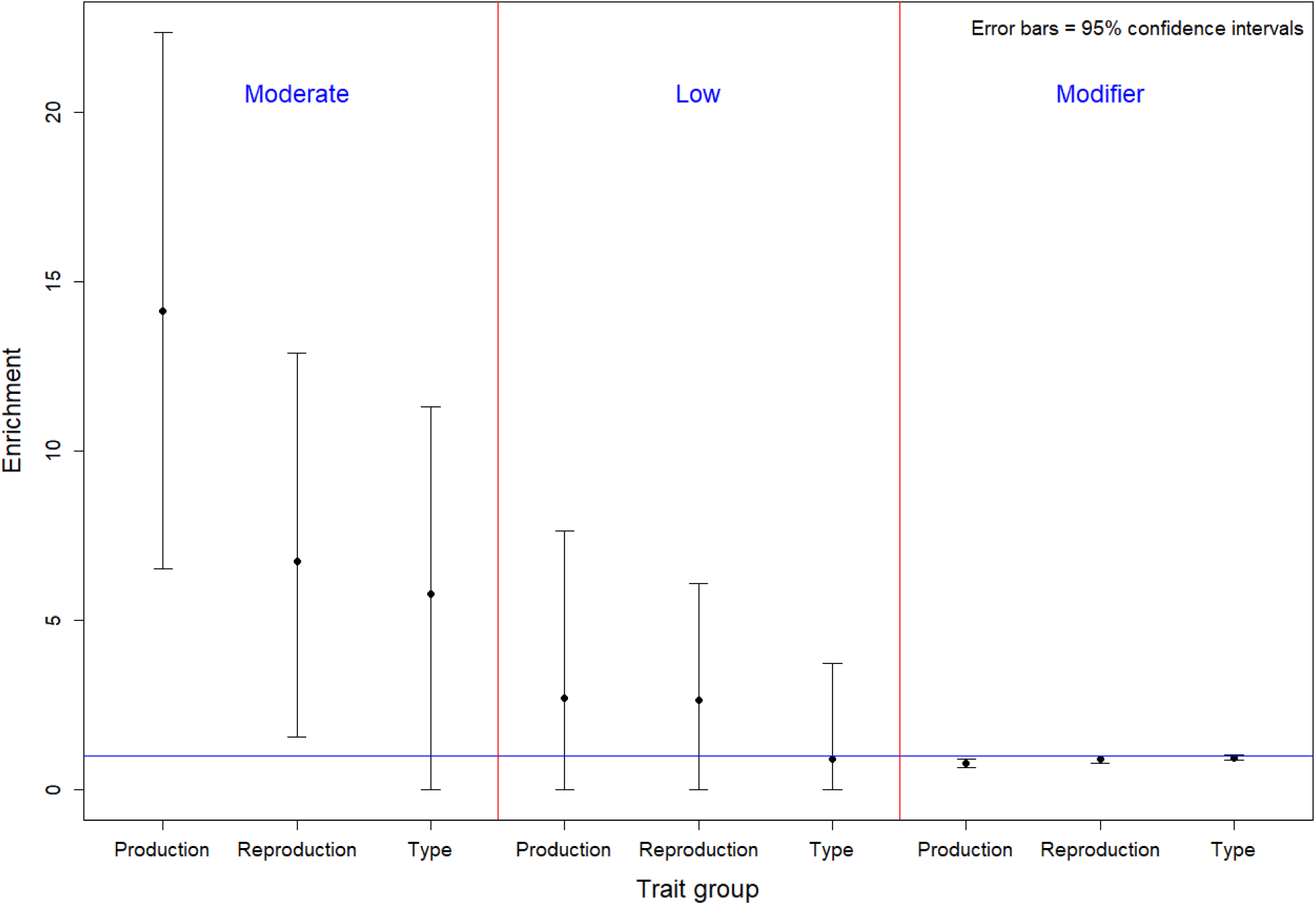
Enrichment estimates for SnpEff predicted impact by three groups of traits.

### Candidate genes

Based on the PPCs of variants after incorporation of SnpEff impact, we calculated PPC for each gene in each independent association signal. In total, there were 564 gene-trait association pairs with PPC >0.01 (Table S22). Most of the genes had either a large (>0.95) or small PPC (<0.05) (Fig S4). We further obtained a short list of the most promising candidates by applying conservative criteria: PPC >0.9 if a gene is associated with only one trait and PPC >0.5 for all traits if a gene affects multiple traits.

This short list had 69 unique genes including both previously reported genes and newly discovered ones for cattle traits (Table 2). For example, *ABCG2* and *DGAT1* are known to affect milk production in dairy cattle ^19,20^. The *ARRDC3* gene has been associated with body confirmation traits and calving traits in beef and dairy cattle ^21–23^. Our fine-mapping study also revealed novel gene/association combinations for dairy traits. A previous study reported that the *ABCC9* gene was associated with fat yield, protein yield, and calving to first service interval in Holstein cattle^24^. In our study, we found a pleiotropic effect of this gene on body type traits (fore udder attachment and udder depth), milk production (milk and protein yields), and daughter pregnancy rate, with a PPC of almost 1 for all the associated traits. In addition, we found that there were no common variants among the credible variant sets for these traits (Table 2), suggesting that *ABCC9* might have different causal mutations for the associated traits. *TMTC2* has been associated with teat length ^23^, and our fine-mapping showed that it had an effect on six type traits (including teat length, fore udder attachment, front teat placement, rear teat placement, rear udder height, and final score), with PPC being ≥0.95 for all those traits. Abo-Ismail et al. reported *CCND2* was associated with stature ^23^. Our fine-mapping results determined its association with four type traits (PPC >0.95 for body depth, rump width, and stature). It is worth noting that our fine-mapping study not only discovered association of a gene with a trait, but also provided the posterior probability of being causal for a gene.

**Table 2.**
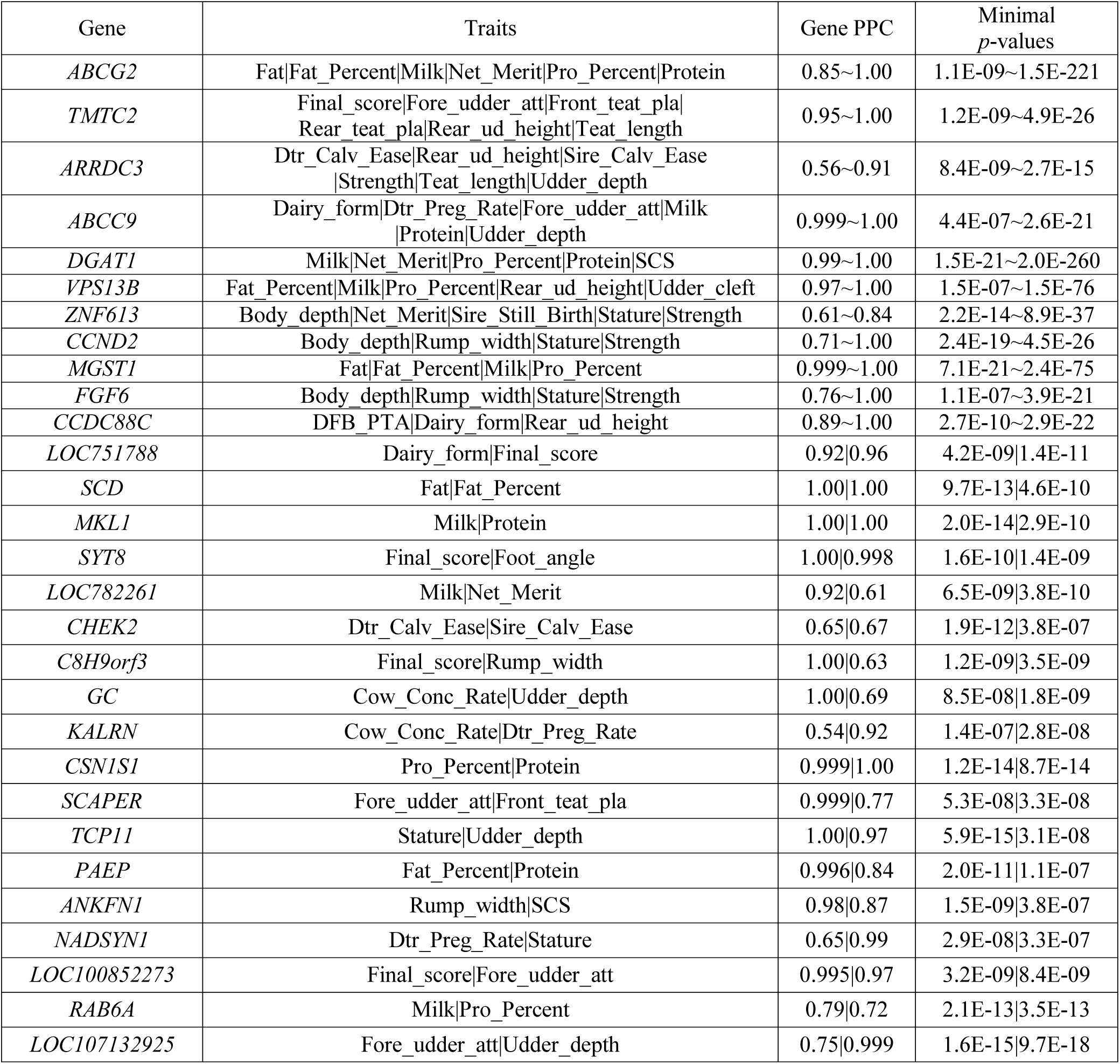
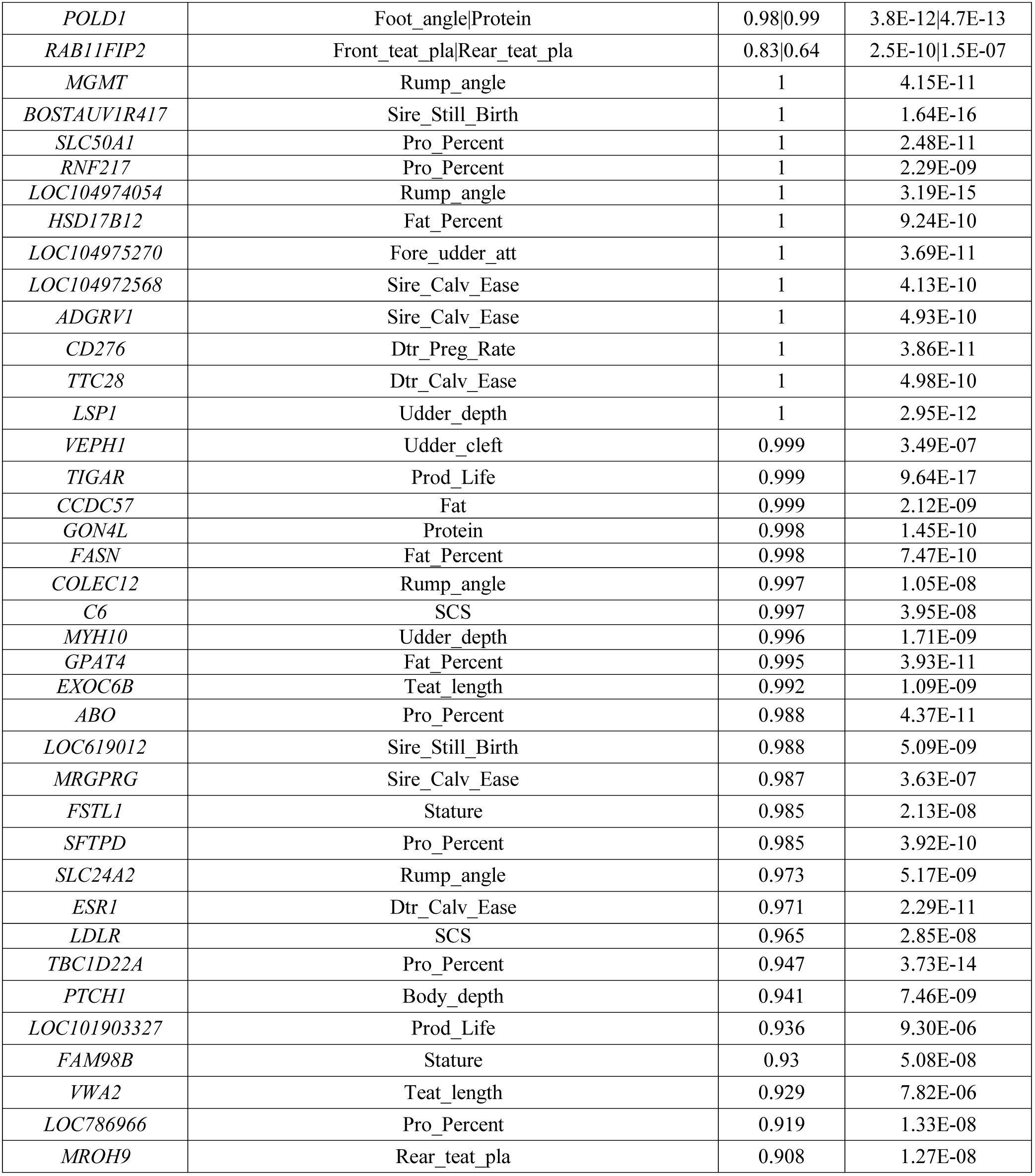
Candidate genes with high posterior probability of causality.

### Candidate variants

Because our stringent quality control filtering during and after imputation removed many variants, fine-mapping of the QTL regions to single-variant resolution could not always be achieved. Nevertheless, we obtained 95% credible variant set for each independent signal and merged them into one table. This resulted in a total of 1,582 unique variants (Table S23). We generated a short list of those variants with a moderate impact on protein coding and PPC >0.2 (Table 3). Among the list, some variants have been previously reported, e.g., Chr6:38027010 in *ABCG2* ^19^ and Chr26:21144708 in *SCD*^25^. We also found other promising candidate variants, e.g., Chr7:93244933 in *ARRDC3* with an average PPC of 0.608 on 9 traits, Chr8:83581466 in *PTH1* with an average PPC of 0.68 on two type traits (body depth and strength), Chr1:69673871 in *KALRN* with an average PPC of 0.46 on two reproduction traits (cow conception rate and daughter pregnancy rate), Chr17:70276788 in *CHEK2* with an average PPC of 0.39 on two calving traits (sire calving ease and daughter calving ease).

**Table 3.** Missense variants with posterior probability of causality >0.2.

| Variant | Gene | MAF | Average PPC | Traits |
| --- | --- | --- | --- | --- |
| 7:93244933 | *ARRDC3* | 0.099 | 0.608 | Body_depth\|Dtr_Calv_Ease\|Net_Merit<br>\|Prod_Life\|Rear_ud_height\|Sire_Calv_Ease<br>\|Strength\|Teat_length\|Udder_depth |
| 6:38027010 | *ABCG2* | 0.015 | 0.87 | Fat\|Fat_Percent\|Milk\|Net_Merit<br>\|Pro_Percent\|Protein |
| 8:83581466 | *PTCH1* | 0.027 | 0.678 | Body_depth\|Strength |
| 26:21144708 | *SCD* | 0.253 | 0.571 | Fat\|Fat_Percent |
| 1:69673871 | *KALRN* | 0.105 | 0.462 | Cow_Conc_Rate\|Dtr_Preg_Rate |
| 19:7521843 | *ANKFN1* | 0.217 | 0.446 | Rump_width\|SCS |
| 29:50290087 | *SYT8* | 0.388 | 0.438 | Final_score\|Foot_angle |
| 29:50286107 | *TNNI2* | 0.203 | 0.436 | Rump_width\|Stature |
| 29:50289940 | *SYT8* | 0.387 | 0.399 | Final_score\|Foot_angle |
| 17:70276788 | *CHEK2* | 0.088 | 0.388 | Dtr_Calv_Ease\|Sire_Calv_Ease |
| 18:57017616 | *POLD1* | 0.103 | 0.291 | Foot_angle\|Protein |
| 8:83044210 | *FANCC* | 0.116 | 0.252 | Rear_teat_pla\|Udder_depth |
| 14:1321450 | *LOC782261* | 0.207 | 0.206 | Milk\|Net_Merit |
| 14:2072259 | *LOC786966* | 0.090 | 0.919 | Pro_Percent |
| 18:44378414 | *CHST8* | 0.120 | 0.889 | DFB_PTA |
| 5:118244695 | *TBC1D22A* | 0.177 | 0.676 | Pro_Percent |
| 5:30259026 | *NCKAP5L* | 0.252 | 0.611 | Teat_length |
| 3:15464749 | *GBA* | 0.063 | 0.601 | Milk |
| 3:20189903 | *ADAMTSL4* | 0.075 | 0.571 | Dairy_form |
| 11:104232298 | *ABO* | 0.309 | 0.449 | Pro_Percent |
| 19:51319797 | *CCDC57* | 0.350 | 0.423 | Fat |
| 18:61020273 | *ZNF331* | 0.038 | 0.322 | Dairy_form |
| 19:51319759 | *CCDC57* | 0.350 | 0.304 | Fat |
| 8:85147150 | *LOC101906801* | 0.117 | 0.302 | Strength |
| 13:58716308 | *C13H20orf85* | 0.116 | 0.297 | Fore_udder_att |
| 11:104232319 | *ABO* | 0.309 | 0.223 | Pro_Percent |
| 14:66328304 | *SPAG1* | 0.119 | 0.222 | SCS |

## Discussion

In this study, we performed GWAS for 35 production, reproduction, and type traits in dairy cattle with a uniquely large data set, and then fine-mapped the GWAS signals to single-gene resolution. With the fast computing method that we developed (BFMAP), we attempted to find causal effects in hundreds of loci each of which contained thousands of variants. We also investigated the functional enrichment patterns of several functional annotation data available in the cattle genome, and incorporated useful functional information into the final fine-mapping. In sum, we provided not only a credible candidate gene list for follow-up functional validation, but also a unique resource that can be easily employed by future functional enrichment studies.

### Single-trait GWAS

In the single-trait GWAS, we found many association signals that have not been discovered (Fig. 1), clearly demonstrating the benefits of using large dairy cattle data for GWAS of complex traits. Reliabilities of de-regressed PTAs were modeled for most of the traits (Table S1). For the traits with small variation of reliability, we observed similar results for the models with and without reliability; e.g., QTLs found when not modeling reliability were largely the same as those by incorporating reliability for fat percentage and daughter pregnancy rate (Fig S5). Interestingly, we observed some deflations in the GWAS of production traits, which could be due to the large QTL effects on these traits including the *DGAT1* gene. Minor inflations were observed in GWAS for calving traits (i.e., calving ease and stillbirth) and final score (Figs S1N-S1R). Although there were many sporadic variants passing the threshold of genome-wide significance (*P* < 5E-8), we could still locate a few credible GWAS peaks where there were a cluster of significant variants.

## Multiple testing in fine-mapping

Initially, our fine-mapping discovered as many as 19 signals in a candidate region for a trait, as we applied a variant inclusion threshold accounting for only the effective number of independent variants (*m*_eff_) at the locus-by-trait level. We also noticed that there were more locus-by-trait association pairs with multiple signals than with only one signal. By examining those with multiple signals, we found the models often contained a strong signal and several much weaker ones. Those weak signals might result from imperfect model fitting of the lead variants in other signals, instead of being true positives. Nevertheless, filtering out these weak signals with genome-wide significance levels did little harm to the discovery of strong ones.

### Enrichment of candidate causal variants

The enrichment results for SnpEff-inferred variant impact in our study were very different from those reported in human studies ^17^. The differences among the four categories in the human study are more distinctive than ours. This is consistent with our anticipation that high LD in cattle genome makes such enrichment difficult to detect. In addition, high-impact variants generally have a lower frequency than other variants and are thus harder to impute in cattle where the number of reference sequences is small and the original genotype data are of moderate density. Nevertheless, we found a considerable enrichment of candidate causal effects in moderate-impact variants. Incorporation of this enrichment into fine-mapping facilitated the discovery of more candidate causal variants (Fig. 5). The discovery of biologically meaningful enrichment patterns will be valuable for the development of new methods to incorporate functional information into fine-mapping and genomic prediction.

### Fine-mapping

Using BFMAP, we pinpointed some promising candidate genes for economically important traits in dairy cattle. It is promising to validate those genes with high posterior probability of causality (Table 2) in future functional studies. In addition, with our new method of functional enrichment analysis in BFMAP, our fine-mapping result of hundreds of QTLs (Table S6) can be readily used to estimate enrichments of causal effects for additional functional annotation data. Thus, we provided an easy-to-use enrichment analysis resource to test the functional annotations that are being generated by the on-going FAANG and related projects for cattle ^12^.

## Methods

### Genotype and phenotype data

Genotype data have been described in more details previously ^5^. Here we provide a brief summary. SNP and insertion-deletion (InDel) calls (sequence variants) from Run 5 of the 1000 Bull Genomes Project ^26^ were released in July 2015. After stringent quality control edits and removal of intergenic and intronic SNPs, 3,148,506 sequence variants were retained for 444 Holstein bulls. The sequence variants and high-density (HD) SNP genotypes of 312K markers for 26,949 progeny-tested Holstein bulls (and 21 Holstein cows) were combined by imputation using the FindHap software (version 3) ^27^. Finally, we had genotypes of 3,148,506 sequence variants for 27,214 Holstein bulls (179 bulls had both sequence and HD genotypes) and 21 cows. Imputation quality from FindHap was assessed with 404 of the sequenced animals as the reference population and 40 randomly selected animals for validation. The sequence genotypes of the validation animals were reduced to HD SNP genotypes and then imputed back to sequence variants. The average imputation accuracy was 96.7% for the 3,148,506 variants ^5^. After excluding HD SNPs, we found an average accuracy of 96.4% for the newly imputed sequence variants. Chromosome-specific imputation accuracy was >95% for all autosomes except Chromosome 12.

All of the 27,214 Holstein bulls used in this study had highly reliable (average reliability >71% across traits) predicted transmitting abilities (PTAs) for 35 production, reproduction, and type traits (Table 1). Transmitting ability is basically the additive genetic values of cattle. Reliability quantifies the amount of information available in a PTA and measures its accuracy ^28^. De-regressed PTAs were used as phenotype in all our analyses, which excludes parent information and reduces the dependence among animals ^29^. Because each of the bulls had many phenotyped daughters, their PTAs were generally of high reliability, even for low-heritability traits (Table 1). The trait definitions are shown in Table 1 and Supplemental Text, Part I. We categorized the 35 traits into three groups, i.e., production, reproduction, and body type, based on a clustering analysis.

### Single-trait GWAS

The software MMAP ^13^ was used for all single-trait GWAS analyses (https://mmap.github.io/). Basically, MMAP efficiently implements a mixed-model approach for association tests that is similar to GEMMA ^30^ but different from EMMAX ^31^; that is, variance component is estimated uniquely for each marker. We used the following model

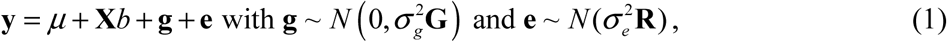

where ***y*** is de-regressed PTAs, *μ* is global mean, ***X*** is genotype of a candidate variant (coded as 0, 1 or 2) and *b* is its effect, ***g*** is a polygenic effect accounting for population structure, and ***e*** is residual. The genomic relationship matrix (***G***) ^32^ was built using ~312K HD SNP markers (filtered by MAF>1%). ***R*** is a diagonal matrix (*R_u_ =* 1/*r*^2^ *−*1), which is used to model differential reliability among animals.

We disregarded variants on the X chromosome. We also filtered out variants with an MAF of <1% or failing Hardy-Weinberg Equilibrium (HWE) test (*p* < 1E-6). After QC, there were ~2.7 million variants to be tested for association. We used a genome-wide significance level of *P* < 5E-8. QTLs were located by finding GWAS peaks where there were a cluster of significant variants. We used a custom Perl script to find all GWAS peaks and further examined each of the peaks based on the Manhattan plots to filter out suspicious ones (i.e., sporadic significant variants). Subsequently, we determined a total of 286 QTLs (Table S2) that were further analyzed in fine-mapping studies.

To find which ones are novel among the 286 QTLs, we compared our result with Cattle QTLdb (release 35 published on April 29, 2018) that contains 113,256 QTLs/associations from 848 publications ^14^. To ensure correct physical positions of QTLs on UMD 3.1, we first extracted the rs identifiers (rs#) of flanking SNPs for each term from the Cattle QTLdb data, and then used the identifiers to find flanking SNPs’ positions on UMD 3.1 in the Ensembl genome variation database. These SNP positions were used as QTL positions. This procedure can rule out QTL terms whose physical positions are inaccurately converted from genetic maps. The Cattle QTLdb release 35 covers 599 different traits, in which we extracted those with the (almost) same definition as the 35 traits in our study (Table S24). For each of the QTLs that we detected, we determined that it is either previously reported if it is within ±500 kb of any QTL/association for the same trait(s) in the Cattle QTLdb or is newly discovered otherwise (Table S2).

### Multi-trait association analysis

Following a previous study ^22^, our multi-trait association tests were based on a chi-square statistic with multiple degrees of freedom. For each variant, the chi-square statistic for the multi-trait association test was calculated by:

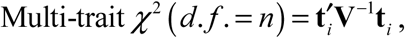

where ***t****_i_* is a *n* × 1 vector of the signed t-values of variant *i* for *n* traits, and ***V*** is an *n × n* correlation matrix for the *n* traits which is calculated using signed t-values of genome-wide variants. In our analysis, the signed t-values were obtained from single-trait GWAS for 2,619,418 variants passing QC, and the correlations between traits were calculated using all the variants. To test the robustness of the estimated correlation using all sequence variants ^33^, we also computed the correlation matrix using two variant subsets obtained by selecting every 10th and every 100th variant. The three variant sets produced similar correlation estimations (Fig S6).

We performed hierarchical clustering based on the absolute correlation coefficients, and then did multi-trait association analysis for each of the three resulting clusters of traits as shown in Fig 2. Specifically, we excluded net merit and days to first breeding (DFB) in production and reproduction clusters, respectively, because these traits are linear combinations of other traits and the number of bulls for DFB was much smaller compared to other traits. We also excluded the four calving traits to avoid sporadic significant variants. Additionally, all the traits except for the six traits aforementioned were analyzed as a whole in a separate multi-trait association test.

### Bayesian Fine-MAPping (BFMAP)

We developed the following Bayesian model for fine-mapping:

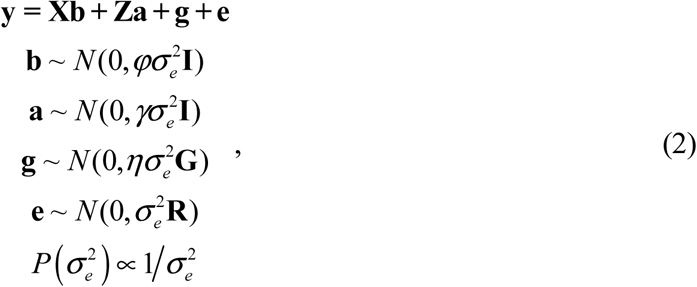

where ***y*** is a phenotype vector of size *n* for a complex trait, ***b*** is a vector of covariate (other than genomic variants) effects and ***X*** is corresponding design matrix, ***a*** is a vector of variant effects and ***Z*** is corresponding genotype coding matrix (e.g., genotype coding for additive, dominance, or imprinting effects ^34^), ***g*** is a vector of polygenic effect for controlling population structure, ***G*** is a corresponding variance structure matrix (e.g., genomic relationship matrix), and ***e*** is the residual with variance structure ***R*** for modelling reliability or accuracy of phenotypic records as in model (1). The common variance component 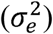 is given by a non-informative Jeffrey’s prior. Other variance parameters (*ϕ*, *γ*, and *η*) are treated as known. Generally, we can set *ϕ* to a large value (e.g., 1E8) to make ***a*** act like fixed effects. A genomic variant is usually considered to have a small but noticeable effect, so we can set *γ* at 0.01 or 0.04 ^35,36^. When ***Za*** only accounts for a tiny proportion of phenotypic variance (this is true when modeling variants from a small genomic region), we can set *η* based on the heritability (*h*^2^) by *η* = *h*^2^ /(1 − *h*^2^). In practice, we can instead use heritability estimate 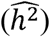 in the null model without variants to determine *η*.

We can easily compute *P(D\M)* (data *D*, and model *M* regarding variant inclusion) by integrating out 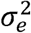 based on model (2). To allow easy calculation, we use a linear transformation to the model (Supplemental Text, Part II). We can further obtain the null distribution of Bayes factors (H_0_: ***a*=0**) in model (2) by an extension of the results by Zhou and Guan ^36^ (Supplemental Text, Part III). Based on the null distribution, scaled Bayes factor ^36^ and corresponding p-value can be computed for our model.

We seek to identify independent association signals within a QTL region and to assign a posterior probability of causality (PPC) to each variant with fine-mapping. Following the method by Huang et al. ^7^, our fine-mapping approach includes three steps: forward selection ^37^ to add independent signals in the model, repositioning signals, and generating credible variant set for each signal. Although our approach uses the same framework as Huang et al. ^7^, there are a few notable differences (Table S25). While they only provided some R scripts for disease data, we provide a fast, general-purpose software tool for fine-mapping analysis of complex traits.

We set *ϕ = γ* = 1*E*8 in model (2) for fine-mapping, which enables easy calculation of *p*-value for a newly added variant conditional on variants already in model (Supplemental Text, Part IV). We use a Bonferroni-corrected threshold ^37^ as stopping criterion in forward selection; that is, forward selection stops when (2logsBF + 1) < 2log *m*_eff_, where *m*_eff_ is the effective number of independent variants calculated using the method by Li and Ji ^38^. Suppose that we select *p* independent signals in forward selection and determine a set of lead variants (*S_1_*) for the *p* signals after repositioning. Then, for signal *i* with lead variant (*I*_i_), we have a variant set (*S_i_*) containing variants that have substantial LD with *I_i_* but weak LD with lead variants in other signals *S_1_* \ {*I*_i_}. Accordingly, we can compute PPC of variant *j* (*vij*) in *S_i_* conditioning on *S_1_* \ {*l*_i_}:

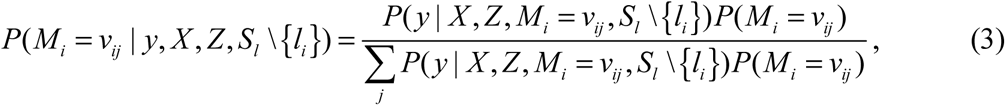

where *M_i_ = v_ij_* denotes that the causal variant in signal *i* is variant *j* in *S_i_* (i.e. *v_ij_*). We can easily get a credible variant set passing a given confidence level (e.g., 95%) for a signal, by sorting variants in a descending order of PPC and including them in the set from top to bottom. We can also calculate PPC of a gene by summing up PPCs of all variants within the gene.

In the study by Huang, et al. ^7^, an equal prior for each variant was used; that is, *P*(*M_i_ = v_ij_*) = 1 ∀*v_ij_* ∈ *S_i_*. Here, we propose a method to apply differential prior probabilities by integrating functional annotation, following a previous study on adjusting significance threshold based on functional annotation in GWAS ^17^. With our fine-mapping procedure, it is usually safe to assume one and only one causal variant in each independent signal. For a functional annotation with several categories, we denote the probability of a causal variant being in category *C* as *p_C_* and the probability of a non-causal variant being of category *C* as *q_C_*. We can accordingly obtain:

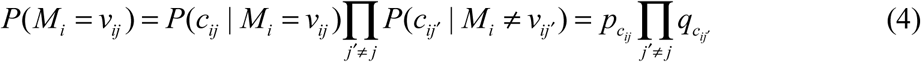

where *c_ij_* denotes the category of variant *j* in *S_i_* (i.e. *v_ij_*).

We estimate *q_C_* with the genome-wide frequencies of the categories ^17^. To estimate *p_C_*, we can use all available independent signals (*M_i_*):

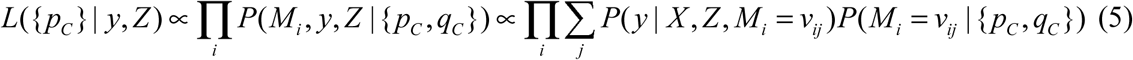

When the signals identified in fine-mapping are independent of each other, we can get:

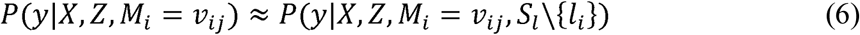

Taking equations (4) and (6) into equation (5), we obtain a likelihood function regarding {*p_C_*} and then get their maximum likelihood estimates (MLEs), 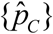. By taking the estimates of {*Pc*, *q_C_*} and equation (4) to equation (3), we get updated PPCs with incorporation of function annotation, which is actually an empirical Bayes approach.

When setting an equal prior for each variant, we find:

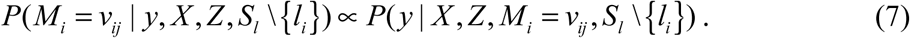

Thus, to estimate {*p_C_*} by equation (5), we can use PPCs from the computation assuming an equal prior for each variant. Accordingly, incorporation of functional annotation includes three steps: computing PPCs given an equal prior for each variant, estimating {*q_C_*} with the genome-wide frequencies of the categories and estimating {*p_C_*} with these PPCs, and updating PPCs with 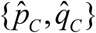. These features make our approach easier to use compared with PAINTOR ^10^ and CAVIARBF ^9^.

### Fine-mapping of dairy cattle traits

Genomic regions for find-mapping were determined by lead variants in single-trait and multi-trait GWAS results. We first determined a minimal region that covered each lead variants (either in single- or multi-trait QTLs), and then extended it 1 Mb upstream and downstream, resulting in a ≥2 Mb candidate region for fine-mapping. The 1-Mb extension allowed the region to cover most variants that have an LD *r*^2^ of >0.3 with the lead variants ^18^.

We obtained a total of 125 loci from single- and multi-trait GWAS results (Table S26). Three loci without enough HD SNPs were removed to ensure imputation quality, thus leaving 122 loci for fine-mapping. A total of 57 loci were associated with more than one trait. Fine-mapping was performed for individual traits, and these 122 loci represented 282 locus-by-trait pairs for 32 traits (three leg traits were excluded for lack of significance). When fine-mapping identified multiple signals in a candidate locus, we kept the strongest one and filtered the rest. The effective number of independent tests was 54,403 for the 282 locus-by-trait pairs (Table S27). Considering that our effective number estimates were already conservative ^39^, we used 5E-7 (<0.05/54,403) as the significance threshold. Subsequently, we found 434 association signals (Table S5).

We found that the locus-by-trait association pairs with more than three signals identified were mostly from still birth and final score (Table S5). We also noticed slight inflation of GWAS results of these two traits (Fig S1). Therefore, we removed the 16 QTLs with >3 fine-mapped signals from all following analyses. We further removed 15 signals whose variant set had ≤10 variants of distinct genotypes, as a small cluster of highly linked variants could indicate inaccurate imputation. Additionally, if there were multiple QTL on a chromosome for a trait, all lead variants in these loci were modeled jointly in fine-mapping. Accordingly, 13 association signals whose lead variant had a *p*-value >5e-7 were removed. After all these edits, we determined a total of 308 association signals (Table S6).

Besides assuming an equal prior for each variant, we further applied differential prior probabilities based on SnpEff-inferred impacts ^15^. Since using equation (5) requires independent association signals, we removed all the association signals for protein, cow conception rate, rear teat placement, udder depth and strength, because they have high correlation (*r*^2^>0.5) with other traits. We also removed another six association signals, since these signals have a substantial LD with another signal (measured by LD *r*^2^ between lead variants >0.25). These edits reduced the number of association signals from 308 to 249. We estimated {*p_C_*, *q_C_*} for variant impact categories based on the 249 association signals, and updated PPCs for all 308 signals by integrating the estimated functional enrichment. Effect impact-incorporated PPCs were used for determining candidate variants or genes. When computing PPC of a gene, all variants within its 2-kb upstream and downstream ranges were included.

### Enrichment analysis in BFMAP

Our enrichment analysis was based on our 249 fine-mapped association signals to estimate *p_C_* (the probability of a causal variant being in category *C*) and *q_C_* (the probability of a non-causal variant being in category *C*). The two probabilities can be estimated using the models described in BFMAP. The enrichment for category *C* is defined as *E_C_* = *p_C_/q_C_* ^17^, for which a value larger than one indicates that candidate causal variants are more enriched in category *C* than across the whole genome. Functional annotations investigated included locations of variants regarding protein-coding genes, effect impact inferred by SnpEff ^15^, and constrained elements predicted by GERP ^16^. Confidence intervals of the enrichment estimates were derived by percentile bootstrap as in ^17^. The association signals were resampled 1,000 times to calculate the confidence intervals. We removed very small categories (like HIGH in SnpEff-inferred effect impacts) in bootstrapping to avoid non-convergence of the maximum likelihood estimation.

### URLs

BFMAP: http://terpconnect.umd.edu/~jiang18/bfmap/
MMAP: https://mmap.github.io/
Cattle constrained elements: ftp://ftp.ensembl.org/pub/release-90/bed/ensembl-compara/68_eutherian_mammals_gerp_constrained_elements/gerp_constrained_elements.bos_taurus.bed.gz
Cattle genome annotation: ftp://ftp.ncbi.nlm.nih.gov/genomes/all/GCF_000003055.6_Bos_taurus_UMD_3.1.1/GCF_000003055.6_Bos_taurus_UMD_3.1.1_genomic.gff.gz
Cattle QTLdb: https://www.animalgenome.org/cgi-bin/QTLdb/BT/index
Cattle genome variation: ftp://ftp.ensembl.org/pub/release-89/variation/gvf/bos_taurus/

### Data availability

The computing program is available at http://terpconnect.umd.edu/~jiang18/bfmap. The original genotype data are owned by third parties and maintained by the Council on Dairy Cattle Breeding (CDCB). A request to CDCB is necessary for getting data access on research, which may be sent to: João Dürr, CDCB Chief Executive Officer. All other data have been shown in the manuscript and supplementary data. All other relevant data are available in the manuscript, Supporting Information files, and from the corresponding author upon request.

## Acknowledgments

We thank the Council on Dairy Cattle Breeding (CDCB) for the access to the genotype data. We thank the 1000 Bull Genomes Project for providing genome references for sequence imputation. We also thank Ellen Freebern for proofreading the manuscript. This work was supported in part by AFRI grant number 2016-67015-24886 from the USDA National Institute of Food and Agriculture (NIFA) and BARD grant number US-4997-17 from the US-Israel Binational Agricultural Research and Development (BARD) Fund. JBC and PMV were also supported by appropriated projects 1265-31000-096-00, “Improving Genetic Predictions in Dairy Animals Using Phenotypic and Genomic Information”, and 8042-31000-104-00, “Enhancing Genetic Merit of Ruminants Through Genome Selection and Analysis”, of the Agricultural Research Service of the United States Department of Agriculture. Mention of trade names or commercial products in this article is solely for the purpose of providing specific information and does not imply recommendation or endorsement by the US Department of Agriculture. The USDA is an equal opportunity provider and employer. The funders had no role in study design, data collection and analysis, decision to publish, or preparation of the manuscript.

## Supporting Information Legends

### Supplemental Text

Part I. Trait definition

Part II. Computation of P(*D*|*M*)

Part III. Null distribution of Bayes factors

Part IV. Bayes factor and *p*-value in fine-mapping

### Supplemental Figures

**Fig S1.** Manhattan plots for 35 dairy traits.

**Fig S2.** Multi-trait association tests compared to single-trait association tests of five production traits on Chr12:56Mb-59Mb. As shown in the panel for protein, the lead variant in protein GWAS (indicated by the left red line) is >1 Mb away from the lead variant in multi-trait association (indicated by the right the red line). Additionally, the lead variant in multi-trait association has a much smaller *p*-value in multi-trait association than in GWAS for any individual trait.

**Fig S3.** Null distribution generated by permutation and actual observation for the ratio of the enrichment using only production traits to the enrichment using reproduction and type traits. Association signals were sampled 1,000 times. Actual observation is denoted by blue lines. *P*-value on the figure was calculated for one tail based on the null distribution. A. The null distribution of the ratio for moderate-impact variants. B. The null distribution of the ratio for low-impact variants. C. The null distribution of the ratio for modifier variants.

**Fig S4.** Histogram of posterior probability of causality for genes.

**Fig S5.** Comparison between GWAS modeling reliability and GWAS not modeling reliability. A. GWAS for fat percentage with reliability modeled. B. GWAS for fat percentage without reliability modeled. C. GWAS for daughter pregnancy rate with reliability modeled. D. GWAS for daughter pregnancy rate without reliability modeled.

**Fig S6.** Estimates of correlations between 35 dairy traits using all sequence variants and two variant subsets. A. Estimates of correlations computed with all sequence variants versus estimates of correlations computed with every 10^th^ variant. B. Estimates of correlations computed with all sequence variants versus estimates of correlations computed with every 100^th^ variant.

### Supplemental Tables

**Table S1.** Genomic control factor and number of QTL of single-trait GWAS for 35 dairy traits.

**Table S2.** QTLs from single-trait GWAS of 35 dairy cattle traits.

**Table S3.** Candidate regions identified by multi-trait association tests compared to those identified by single-trait GWAS.

**Table S4.** The list of candidate regions identified by multi-trait association tests but missed by single-trait GWAS.

**Table S5.** Number of fine-mapped signals for each trait-region association pair after filtering by a significance threshold of 5*E*-7.

**Table S6.** All fine-mapping results.

**Table S7.** Enrichment estimates for location of variants regarding protein-coding genes.

**Table S8.** Enrichment estimates for SnpEff effect impact.

**Table S9.** Enrichment estimates for GERP constrained elements.

**Table S10.** Enrichment estimates for effect impact by GERP constrained elements.

**Table S11.** Enrichment estimates for effect impact generated using three groups of traits.

**Table S12.** All genes fine-mapped with posterior probability of causality of >0.01.

**Table S13.** Variants merged from 95% credible variant sets of all fine-mapped association signals.

**Table S14.** QTLdb traits corresponding to the 35 traits in this study.

**Table S15.** Differences between BFMAP and the fine-mapping approach by Huang et al.

**Table S16.** Genomic regions for fine-mapping.

**Table S17.** Number of variants and effective number of independent variants in genomic regions for fine-mapping.

